# Multi-segment mouse-adaptation of a recent B/Victoria-lineage virus independent from hemagglutinin and neuraminidase

**DOI:** 10.1101/2025.10.12.681926

**Authors:** Arne Matthys, Laura Amelinck, Anouk Smet, Tine Ysenbaert, Thorsten U. Vogel, Xavier Saelens, João Paulo Portela Catani

## Abstract

Influenza B viruses (IBVs) contribute significantly to the annual influenza epidemics in human. Most IBV strains are non- or poorly pathogenic in mice, which are frequently used for vaccine studies. We describe the generation of a mouse-adapted IBV strain that retains pathogenicity in mice when carrying hemagglutinin (HA) and neuraminidase (NA) gene segments from a heterologous IBV strain. Serial passage of an influenza B reassortant virus, containing the HA and NA segments from B/Washington/02/2019 on a mouse-adapted B/Memphis/12/1997 backbone, resulted in the selection of an IBV that was highly pathogenic for mice. This mouse-adapted IBV strain had acquired non-synonymous mutations in 5 gene segments. Sequence analysis of the intermediate passages indicated that mutations in the matrix (M), polymerase acidic (PA), and polymerase basic 1 (PB1) gene segments appeared at passages 9 and 13, suggesting that these mutations contributed to the pathogenicity in mice. Mouse challenge studies with rescued reassortant viruses with one or multiple mutated gene segments, confirmed the importance of substitutions in the M and PA segments for pathogenicity. Using the novel mouse-adapted IBV backbone, we rescued reassortant viruses containing the HA and NA segments of B/Austria/1359417/2021 and demonstrated its increased pathogenicity in BALB/c mice compared to IBV rescued on the parental strain. This mouse-adapted IBV backbone provides a valuable tool for the study of IBV in mice.

## Introduction

Influenza B viruses (IBVs) are human respiratory pathogens that cause a significant yearly recurrent disease burden (1). IBVs diverged in the 1980s into two antigenically distinct lineages, named the B/Victoria and B/Yamagata lineage. These lineages have been circulating with alternating predominance (2). Since 2020, however, B/Yamagata lineage IBVs have not been detected. Given that IBVs lack an established animal reservoir, the continued absence of detectable B/Yamagata lineage viruses suggest their extinction (Caini et al., 2024), (Koutsakos et al., 2024).

The segmented, negative-sense RNA genome of IBVs is replicated by the viral RNA-dependent RNA polymerase that lacks proofreading activity, contributing to the genetic variability. Nucleotide substitutions occur in all gene segments and are generally higher for B/Victoria compared to B/Yamagata lineage viruses (4). With 2×10^−3^ substitutions/site/year, the IBV hemagglutinin (HA) gene mutates generally slower than the HA of human H1N1 (4×10^−3^ substitutions/site/year) and H3N2 (5.5×10^−3^ substitutions/site/year) influenza A viruses (5). Despite the slightly lower substitution rate, antigenic drift in IBV necessitated frequent updates of the recommended vaccine strains (e.g. 6 distinct IBV strains in the last 10 years) (6).

IBVs are poorly pathogenic to mice, limiting the use of this small animal model for IBV research. Serial passage of IBV through mouse lungs is a well-established method to select a pathogenic virus. IBV adaptation to mice has been associated with amino acid substitutions across all eight viral gene segments (7–10). Identification of the acquired mutations combined with reverse genetics allows to identify the substitutions that are necessary and sufficient for the mouse-adapted phenotype. Mouse-adapted IBVs have been used to define pathogenic determinants and were exploited for the generation of live attenuated influenza vaccines candidates (11), to study the antibody repertoire and antigenic drift (12), or for drug susceptibility testing (13).

In this study, we report the serial passage of an IBV through mice that resulted in the establishment of a backbone of internal segments that conferred increased mice pathogenicity of IBV reassortants.

## Results

The HA and neuraminidase (NA) segments of B/Washington/02/2019 were used to rescue a 2:6 reassortant virus using the internal segments obtained from the mouse-adapted B/Memphis/12/1997. This initial backbone includes the single substitution (N221S) in BM1, previously shown to confer increased pathogenicity to B/Memphis/12/1997 (10). Sequence analysis revealed that the rescued reassortant IBV, which was named B/HwNw-Mem97, carried an additional T211A substitution in the HA and M390I in the NA segment. Despite encoding the N221S substitution in the C-terminus of BM1, previously described to enhance mouse virulence, the B/HwNw-Mem97 reassortant virus was not pathogenic to BALB/c mice (Figure 1a). Similarly, B/HwNw-Mem97 was not pathogenic to DBA/2J mice, a mouse strain characterized by increased sensitivity to influenza virus infection **(Figure 1b)** (14,15).

**Figure 1.**
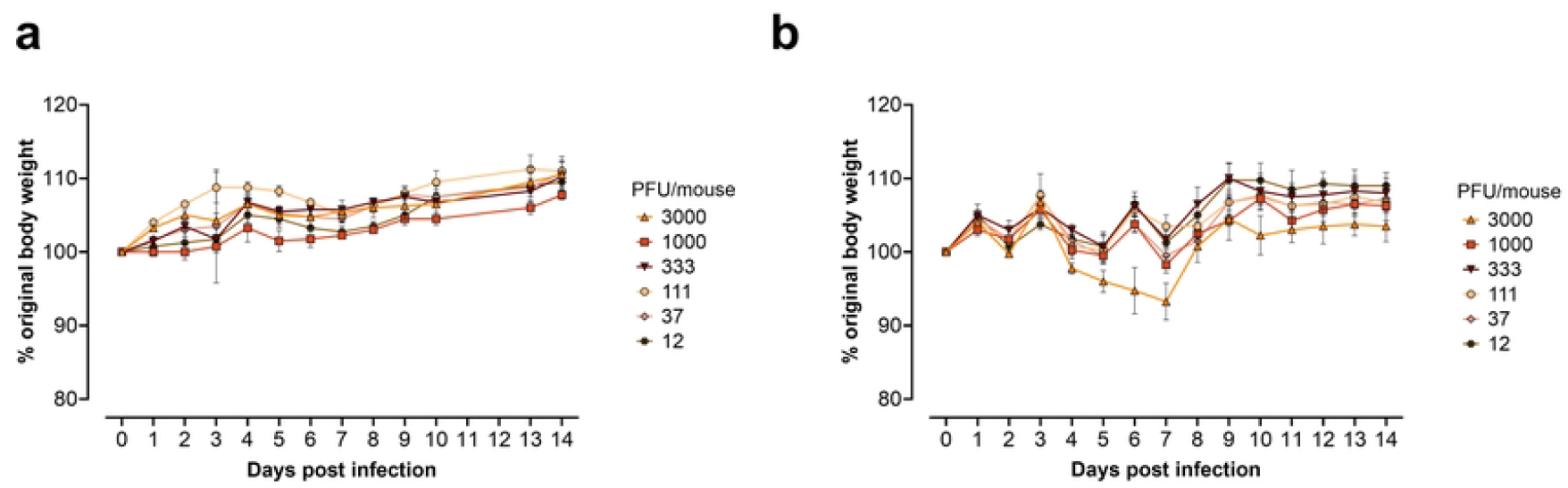
Weight loss induced by B/HwNw-Mem97. The 2:6 B/HwNw-Mem97 reassortant with N221S substitution in the C-terminus of BM1 was inoculated with different amounts in groups of four BALB/c (a) or DBA/2J mice (b). Data points represent the mean relative body weight and error bars the standard error of the mean.

To isolate an IBV strain with enhanced pathogenicity, the B/HwNw-Mem97 virus was passaged multiple times through the lungs of DBA/2J mice and propagated on MDCK cells between each *in vivo* passage. After 13 passages, the infected mice showed clinical signs of disease (ruffled fur and reduced mobility). The virus that was obtained after passage 13, referred to as the B/HwNw-Mem97 m.a. virus, was amplified on MDCK cells and titrated in BALB/c and DBA/2J mice. The resulting 50% lethal dose (LD_50_) were 810 PFU for BALB/c and 25 PFU for DBA/2J mice **(Figure 2)**.

**Figure 2.**
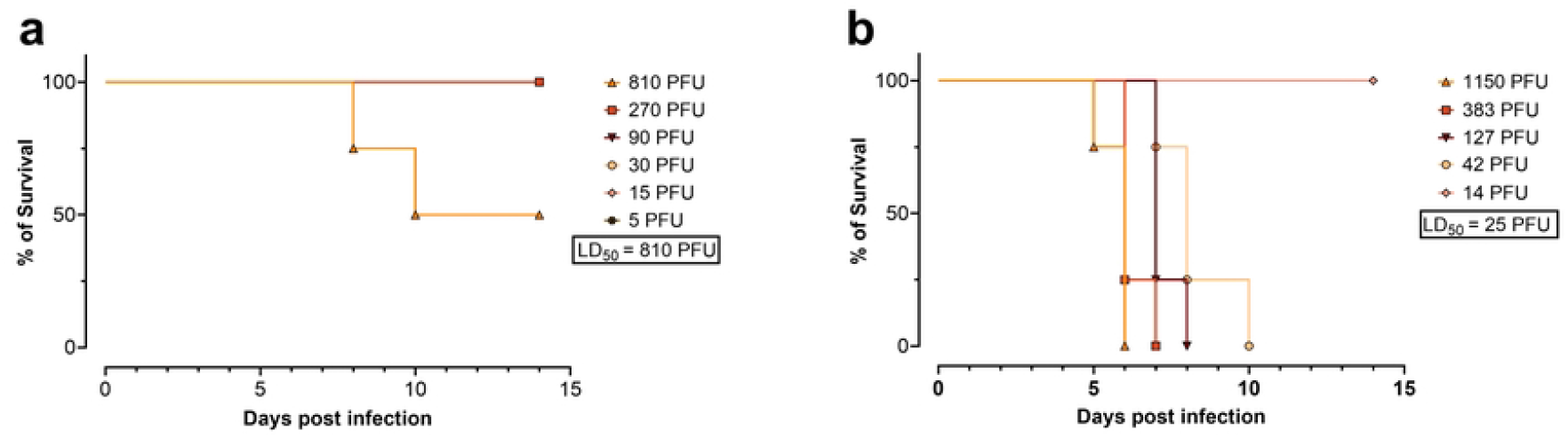
Pathogenicity of B/HwNw-Mem97 m.a. virus. BALB/c (**a**) and DBA/2J mice (**b**) were inoculated with different doses of B/HwNw-Mem97 m.a. virus (obtained after thirteen serial passages in DBA/2J mice). The LD_50_ was calculated using the Reed and Muench method.

To identify the mutations associated with increased pathogenicity, the complete B/HwNw-Mem97 m.a. genome was sequenced. Three synonymous mutations were identified, one in the PB2, one in the PB1, and one in the BM1 segment. In addition, a total of eight non-synonymous mutations were found across PB2, PB1, PA, NA, and BM1 gene segments (**Figure 3a, Table 1**). To understand the evolutionary trajectory of each mutation, the segments of intermediate viral passages 3, 6, and 9 were subcloned in the pHW2000 vector and sequenced (**Figure 3b, Table 1**).

**Table 1.**
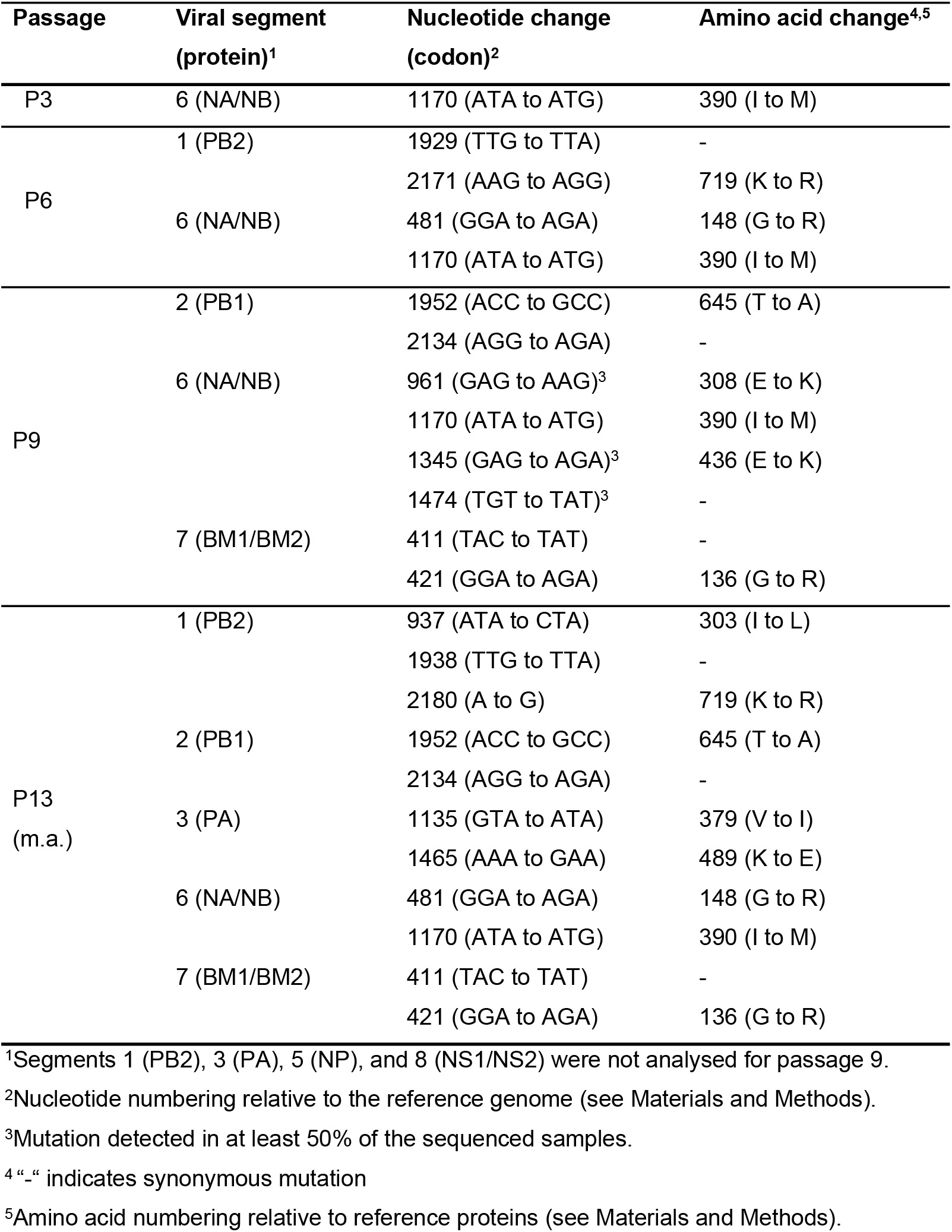
Mutations present in the B/HwNw-Mem97 m.a. genome relative to the parental rescued virus. Mutations were identified in intermediate passages (P3, P6, or P9) with Sanger sequencing and in the mouse adapted virus (P13) with NGS and Sanger sequencing.

**Figure 3.**
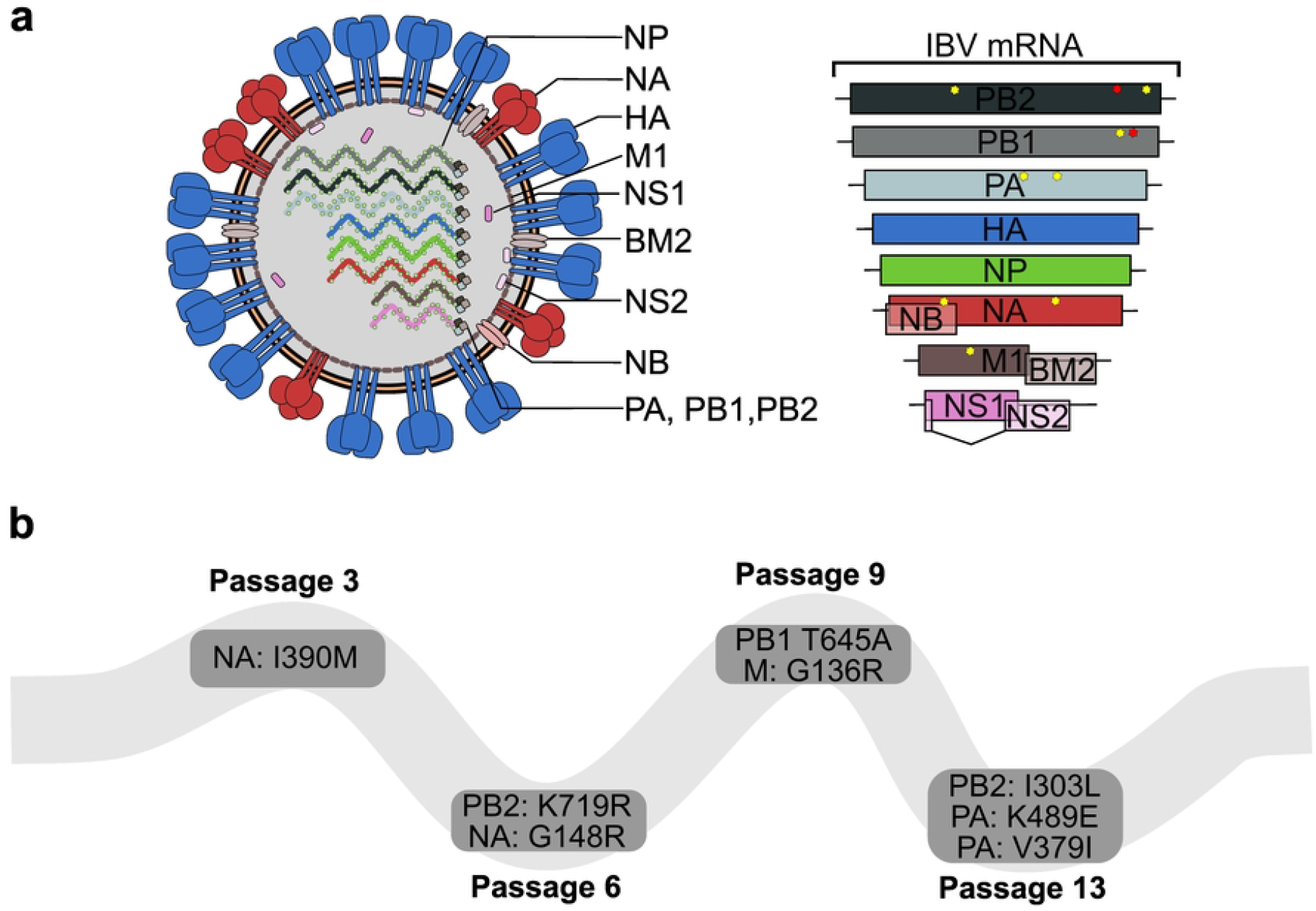
Multi-segment mouse-adaptation of the B/HwNw-Mem97 m.a. virus. **a)** Eight gene segments code for at least 11 IBV proteins. Nucleotide substitutions in IBV after 13 in vivo passages are mapped per segment with yellow or red stars for non-synonymous and synonymous mutations, respectively. **b)** Appearance of mutations resulting in amino acid substitutions present in the B/HwNw-Mem97 m.a. virus, through the tested passages.

Next, reassortant viruses were rescued using combinations of segments from B/HwNw-Mem97 m.a. virus to investigate the individual contributions of each segment to the pathogenicity. These reassortants consisted of combinations of segments derived from the parental B/HwNw-Mem97- and the B/HwNw-Mem97 m.a. virus (**Figure 4a**). Despite multiple attempts, we were unable to rescue B/HwNw-Mem97 m.a. by reverse genetics. We inoculated BALB/c mice with an equal amount of the different reassortant viruses (1620 PFU/mouse, which corresponds to 2 LD_50_ of B/HwNw-Mem97 m.a.) and monitored morbidity and mortality for 14 days (**Figure 4b**). All mutated segments except PB2, or the combination of PB2 and PB1, contributed significantly to morbidity. Mortality was only significantly affected by the combination of mutated segments PB1 and PA, or NA and M. PA and M segments derived from B/HwNw-Mem97 m.a. were predominantly and independently associated with increased mortality. Infection of BALB/c mice with reverse genetics B/HwNw-Mem97 viruses with HA and PB2 segments derived from B/HwNw-Mem97 m.a. or B/HwNw-Mem97 with HA, PB2 and PB1 segments derived from B/HwNw-Mem97 m.a., resulted in limited body weight loss (**Figure 4b**).

**Figure 4.**
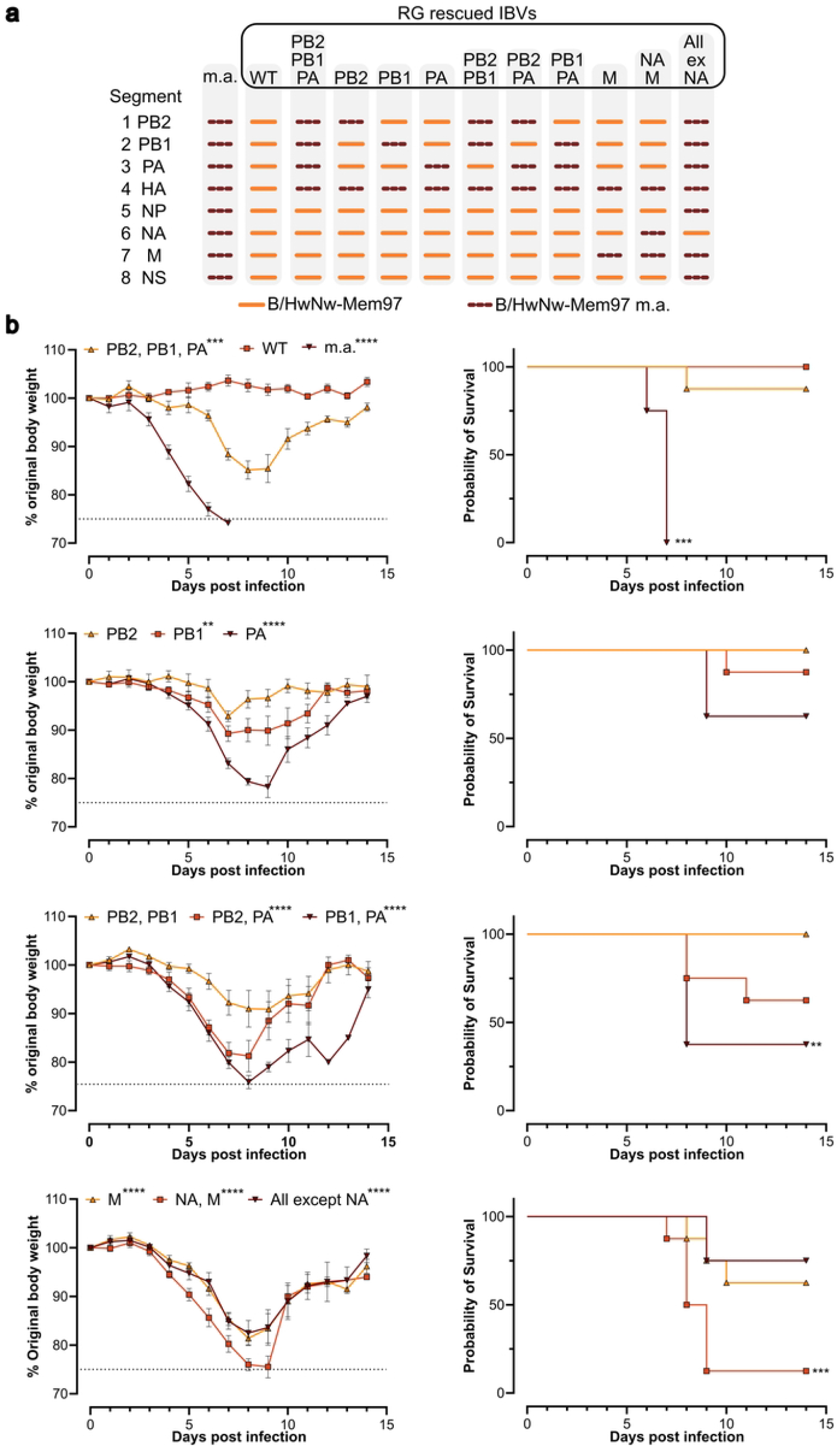
Morbidity and mortality in BALB/c mice challenged with different B/HwNw-Mem97 m.a. reassortant viruses. **a)** Genomic composition of the B/HwNw-Mem97 m.a. (m.a.) and rescued reverse genetics (RG) IBV viruses. Note that the sequences of the HA, NP, and NS segments of B/HwNw-Mem97 and B/HwNw-Mem97 m.a. are identical. **b)** Bodyweight (left) and survival curves (right) of BALB/c mice over time following challenge with 1620 PFU/mouse. Differences in bodyweight between groups (compared to the WT group, shown only in the first graph) were tested by two-way ANOVA with Dunnett’s multiple comparison: **p=0.0021, ***P<0.0002, ****P<0.0001. Differences in survival (compared to the WT group) were tested with a log-rank (Mantel-Cox) test: **p=0.0021, ***p<0.0002, ****p<0.0001. The graphs show compiled data from two independent experiments (n = 4 mice per group per experiment). The dotted lines in the bodyweight graphs indicate the ethical maximum permitted bodyweight reduction.

To assess whether the 6 internal gene segments of B/HwNw-Mem97 m.a. could confer pathogenicity in mice when combined with heterologous HA and NA, we rescued recombinant viruses carrying the HA and NA gene segments from B/Austria/1359417/2021 and either the six internal segments from B/HwNw-Mem97 m.a. or from B/Memphis/12/1997 m.a. (with the BM1 N221S substitution; (10)). BALB/c mice were inoculated with 450 PFUs of each reassortant. Although mice in both groups experienced body weight loss and mortality, the reassortant virus bearing the B/HwNw-Mem97 m.a. internal genes induced significantly greater weight loss. The mortality outcome did not differ significantly (**Figure 5**).

**Figure 5.**
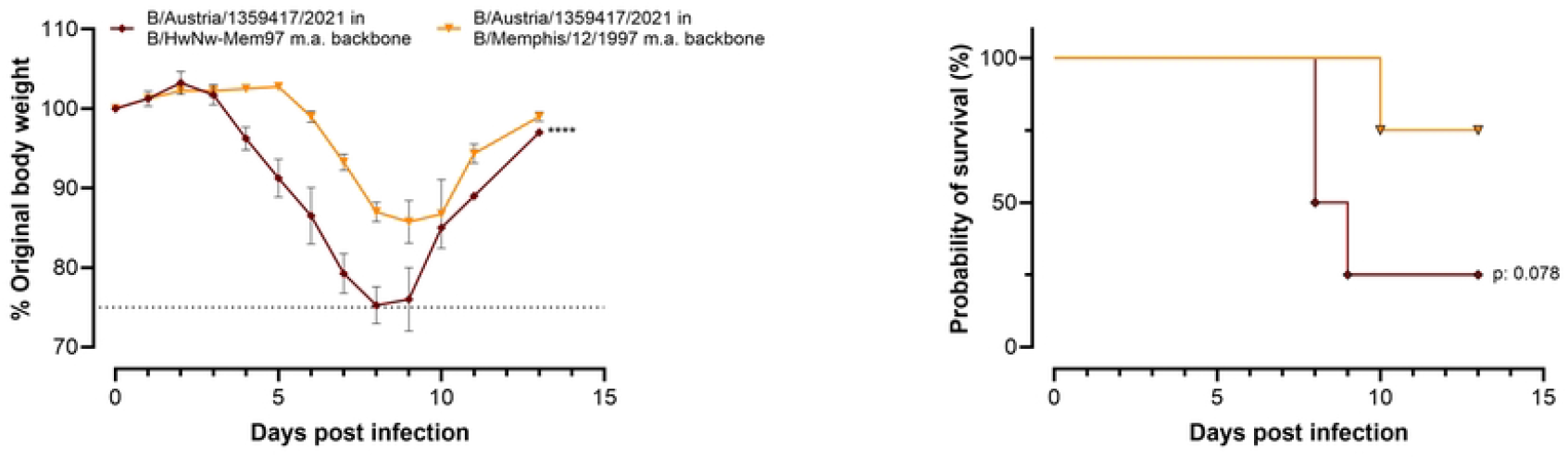
The internal gene segments of B/HwNw-Mem97 m.a. confer increased pathogenicity of RG B/Austria/1359417/2021 virus to BALB/c mice. Bodyweight and survival curves of female BALB/c mice over time following challenge with 450 PFU/mouse (n=4 per group). Differences in bodyweight between groups were tested by two-way ANOVA with Dunnett’s multiple comparison: ****p<0.0001. Differences in survival were tested with a log-rank (Mantel-Cox). The dotted lines in the bodyweight graphs indicate the ethical maximum permitted bodyweight reduction.

## Discussion

Aiming to generate an IBV challenge model bearing the HA and NA components of the B/Washington/02/2019 virus, we rescued a 2:6 reassortant strain, which we named B/HwNw-Mem97, with the internal segments of the mouse adapted B/Memphis/12/1997 (10). The rescued RG B/HwNw-Mem97 was not pathogenic in BALB/c and DBA/2J mice. To generate a pathogenic strain, 13 serial passages were performed in lungs of DBA/2J mice. The resulting mouse adapted IBV, named B/HwNw-Mem97 m.a., had acquired non-synonymous mutations in the PB2, PB1, PA, NA and BM. Among these, the substitutions in the M and PA segments played a predominant role in increasing mice pathogenicity.

A substitution at the C-terminus of the M segment (N221S) has previously been described as a determinant of IBV mouse pathogenicity (10). Although this substitution was present in our rescued B/HwNw-Mem97 virus, this virus did not cause detectable morbidity in mice. After 9 serial passages, the substitution G136R was identified in the M segment and was found to correlate with increased mouse pathogenicity. Substitutions in the M segment have also been linked to increased pathogenicity in mice of influenza A viruses, however the mechanism remains unclear and may involve altered vRNP binding (16).

In the PA segment, a single substitution (K338R) has been reported to enhance pathogenicity in mice by approximately 10-fold across both IBV lineages, and this effect was associated with increased replicase activity (9). However, pathogenicity still required relatively high viral doses (LD_50_ of 10^3.5^ PFU and 10^5.5^ PFU for B/Victoria- and B/Yamagata-lineage viruses, respectively). The B/HwNw-Mem97 m.a. PA bears two substitutions (V379I and K489E). Interestingly, high-yielding egg-adapted IBV backbones were recently described and include the PA substitution K498E (17).

Adaptation of influenza viruses to mice can result in substitutions in HA and NA, which may be undesirable when studying immune responses targeting these antigens. For example, seventeen serial passages in mice of a B/Victoria clade 1A virus (B/Novosibirsk/40/2017) resulted in substitutions in HA (T214I) and NA (D432N) (8). Seventeen passages of B/Florida/04/2006 resulted in 5 amino acid changes, of which one in HA (D424G) (18).

The B/HwNw-Mem97 m.a. virus as reported here carries the HA substitution T211A and NA substitutions G148R and I390M. Notable, the T211A in HA and M390I in NA were present in the rescued parental strain (P0), and consequently occurred independently of the mouse adaptation. The methionine at position 390 in NA is highly conserved among IBVs. Although the rescued B/HwNw-Mem97 m.a. initially carried this M390I substitution, it was reverted back to a methionine (I390M) after, at the most, three passages. Thus, given that the NA segment at least partially contributed to pathogenicity in mice, it is likely that the G148R substitution in NA is associated with increased pathogenicity. Typically, position 148 in NA features a glycine in B/Victoria-lineage viruses or a glutamic acid in B/Yamagata-lineage viruses. Neighbouring residues (especially D149) are critical components of the catalytic site, directly participating in sialic acid receptor binding (19). Substitution G148R may introduce local allosteric changes that could have a limited contribution to viral pathogenicity, since reassortant viruses with the parental NA segment were also pathogenic in mice.

The rescue of the 2:6 B/Austria/1359417/2021 reassortant on our B/HwNw-Mem97 m.a. backbone resulted in a virus with increased pathogenicity in comparison to its counterpart rescued on the B/Memphis/12/1997 mouse adapted backbone. No substitutions were found in the HA or NA derived from B/Austria/1359417/2021 in the reassortant virus, indicating that mouse pathogenicity relied solely on the m.a. internal segments, allowing the study in mice of IBV viruses with potentially unaltered antigenicity.

In summary, we isolated an IBV backbone carrying non-synonymous substitution in 5 gene segments (PB2, PB1, PA, NA, and M). Except for PB2, all the mutated segments seem to, at least partially, contribute to increased mice pathogenicity. The m.a. backbone was successfully used to rescue a second virus comprising HA and NA of the recent B/Austria/1359417/2021 vaccine strain. The described backbone offers a new platform to rescue IBV strains that are pathogenic in mice, without altering their surface antigenicity.

## Materials and methods

### Virus rescue

HA and NA sequences from B/Washington/02/2019 (GISAID accession numbers EPI1368874 and EPI1368872, respectively) and B/Austria/13595417/2021 (GISAID accession numbers EPI1845793 and EPI1845794, respectively) were generated by custom gene synthesis (GeneArt, Thermo Fischer) and cloned in the bidirectional expression plasmid pHW2000. The gene segments from the mouse adapted B/Memphis/12/1997 (10) were reverse transcribed from the virus stock, using the primers described previously (20) and cloned in the pHW2000 expression vector. To rescue reassortant viruses, a co-culture of MDCK and HEK293 cells was transfected with 1 μg of plasmids encoding HA and NA of B/Washington/02/2019, and the 6 plasmids (PB2, PB1, PA, NP M, and NS) encoding internal proteins of the mouse adapted B/Memphis/12/1997. The supernatant (SN) was recovered after observation of cytopathic effect and titrated (PFU determination). All rescued viruses were amplified on MDCK cells to prepare working stocks. Briefly, MDCK cells were inoculated with a MOI of 0.001 in the presence of TPCK-treated trypsin (2 μg/mL, Sigma T1426). After cytopathic effect was evident, the cell culture supernatant was recovered, cleared from cells and debris by centrifugation, and titrated by plaque assay on MDCK cells (PFU determination).

### Mouse adaptations

The initial viral infection was performed with 1 x 10^5^ PFU (50 µL per nostril). During each passage, groups of 3 DBA/2J mice were inoculated with a 1/100 dilution of the viral stock and euthanized 72 hr later to recover virus from the lungs. Lung homogenates were prepared in PBS using sterile metal beads and a Tissue Lyser II mechanical homogenizer. The homogenates were clarified by centrifugation at 1,000 × g for 10 minutes at 4°C, then pooled and diluted 1:1000 prior to inoculation of MDCK cells. Infected cells were maintained in the presence of 2 µg/mL TPCK-treated trypsin (Sigma-Aldrich, T1426) until observation of cytopathic effect. The cell culture medium was then recovered, cleared by centrifugation and used at 1/100 dilution to inoculate naive mice. These passages were repeated 13 times when clinical signs of influenza virus infection in mice (ruffled fur and reduced mobility) was observed on day 3 after inoculation.

### Virus sequence analysis

The mouse-adapted and parental strains were sequenced using Oxford nanopore at PathoSense (Belgium). The internal genes of intermediate passages were reverse transcribed using the primers described by (21), and the amplified segments were cloned into the bidirectional plasmid pHW2000 and sequenced by Sanger sequencing (IDT Belgium). Nucleotide substitutions are numbered relative to the B/Washington/02/2019 reference genome retrieved from the NCBI viral genomes resource database (22) with the following accession codes for HA (MK676294) and NA (MK676296). Nucleotide substitutions in the internal gene segments are numbered relative to the B/Memphis/12/1997 reference genome with following accession codes for PB1 (AY260942), PB2 (AY260943), PA (AY260944), NP (AY260946), M (AY260941), and NS (AY260948).

### Mouse housing and viral challenge

Specific pathogen-free female BALB/c and DBA/2J mice were obtained from Janvier (France). The animals were housed in a temperature-controlled environment with 12 h light/dark cycles; food and water were provided ad libitum. The animal facility operates under the Flemish Government License Number LA1400563. All experiments were done under conditions specified by law and authorized by the Institutional Ethical Committee on Experimental Animals (Ethical Application EC2023-120 and EC2023-140). Intranasal administration of virus dilution (50 µL per nostril) was performed under isoflurane sedation in a BSL2 graded facility. The ethical endpoint for euthanasia of the mice was reached when mice had lost 25% or more of their body weight relative to the day of challenge. Mice were humanely euthanized by cervical dislocation. The LD_50_ was calculated using the Reed and Muench method (23).

## Acknowledgments

The mouse adapted B/Memphis/12/1997 was kindly provided by Dr. Jonathan McCullers and the pHW2000 plasmid from Dr. Erich Hoffmann and Dr. Robert Webster (St. Jude Children’s Research Hospital, Memphis, TN, USA). We are grateful to the animal caretakers of the animal house at the VIB-UGent Center for Inflammation Research.

## Author Contributions

Conceptualization: João Paulo Portela Catani, Thorsten U Vogel, Xavier Saelens

Methodology: Arne Matthys, Xavier Saelens, João Paulo Portela Catani

Investigation: Arne Matthys, Laura Amelinck, Anouk Smet, Tine Ysenbaert, João Paulo Portela Catani

Formal analysis: Arne Matthys, Laura Amelinck, Anouk Smet, Tine Ysenbaert, João Paulo Portela Catani

Funding acquisition: Xavier Saelens, Thorsten U Vogel Supervision: Xavier Saelens, João Paulo Portela Catani

Writing – original draft: Arne Matthys, João Paulo Portela Catani Writing – review and editing: Thorsten U Vogel, Xavier Saelens

